# Microphase separation of living cells

**DOI:** 10.1101/2022.05.25.493184

**Authors:** A. Carrère, J. d’Alessandro, O. Cochet-Escartin, J. Hesnard, N. Ghazi, C. Rivière, C. Anjard, F. Detcheverry, J.-P. Rieu

**Affiliations:** University of Lyon, Université Claude Bernard Lyon 1, CNRS, Institut Lumière Matière, F-69622, VILLEURBANNE, France

## Abstract

Self-organization of cells is central to a variety of biological systems and physical concepts of condensed matter have proven instrumental in deciphering some of their properties. Here we show that microphase separation, long studied in polymeric materials and other inert systems, has a natural counterpart in living cells. When placed below a millimetric film of liquid nutritive medium, a quasi two-dimensional, high-density population of *Dictyostelium discoideum* cells spontaneously assemble into compact domains. Their typical size of 100 μm is governed by a balance between competing interactions: an adhesion acting as a short-range attraction and promoting aggregation, and an effective long-range repulsion stemming from aerotaxis in near anoxic condition. Experimental data, a simple model and cell-based simulations all support this scenario. Our findings establish a generic mechanism for self-organization of living cells and highlight oxygen regulation as an emergent organizing principle for biological matter.

---

It is not uncommon for physical concepts to migrate to the biological realm [1, 2]. Even though cells are governed by specific genetic programming, their collective behavior share some common generic features with non-living systems. States of matter, such as solid, liquid, gas phases originally defined for molecular components, are now employed to characterize assemblies of cells [3]. Extended analogies with liquids have illuminated the properties of tissues, whose surface tension induces cell sorting, shapes of minimal area, spreading and dewetting [1, 4, 5]. A variety of out-of-equilibrium processes have also proven useful: examples range from the glass and jamming transitions for epithelia [6–8] to the directional solidification model for the growing yeast dynamics [9] or the diffusion-limited aggregation for fractal-like bacterial colonies [10]. Over the years, soft condensed matter ideas have found fruitful applications in cell assemblies.

One soft matter phenomenon that has attracted considerable attention for half a century is micro-phase separation, a process leading to spontaneous formation of equilibrium domains with finite length scale. In the most prominent instance of diblock copolymers, separation is driven by the repulsion between two chemically different blocks but is counteracted by chain connectivity and entropy [11]. The balance between those competing trends results in ordered morphologies with a thermodynamically preferred domain size [12, 13]. The microphase separation of polymers is in fact representative of a generic scenario for pattern formation: modulated phases induced by competing interactions, usually a short-range attraction opposed by a long-range repulsion [14]. This mechanism was recognized in several physical systems, including Langmuir monolayers [15], magnetic films [16], superconductors [17], liquid crystals [18] and clusters of proteins or colloids [19, 20] A related class of phenomena recently uncovered is the formation of biomolecular condensates, whether micron-scale compartments in eukaryotic cells [21] or euchromatin domains in the nucleus [22]. All instances of microphase separation identified so far involve subcellular processes or inert matter. To our knowledge, it has never been observed in populations of cells.

We report here how living cells can self-organize in reversible finite-size domains, a bona fide analogue of the microphase separation in inert matter. Cell-cell adhesion acts as a short-range attraction promoting aggregation but is counteracted by oxygen depletion and aerotaxis, whose combination induces an effective long-range repulsion. As a consequence, the size domain is set by a delicate balance between cell density and oxygen availability. An analytical model and microscopic cell simulations both corroborate this simple picture. Our findings add a new piece to the mounting evidence that oxygen regulation may govern spatial patterning in various contexts, from morphogenesis of eukaryotes to tumor escape and growth [23–25].

## Experiments

Cells of *Dictyostelium Discoideum* (*Dd*) were seeded on a substrate submerged with a film of nutrient culture medium (Fig. 1A-D) and imaged by transmission microscopy for several days (SM-A). As their number increases through division, their spatial distribution becomes heterogeneous: cells gather in aggregates – locally denser regions – which grow by addition from the surroundings or merging with neighboring aggregates (Fig. 1B-C and movie M1). The steady state eventually reached (SM-B) is the focus of this work.

**FIG. 1.**
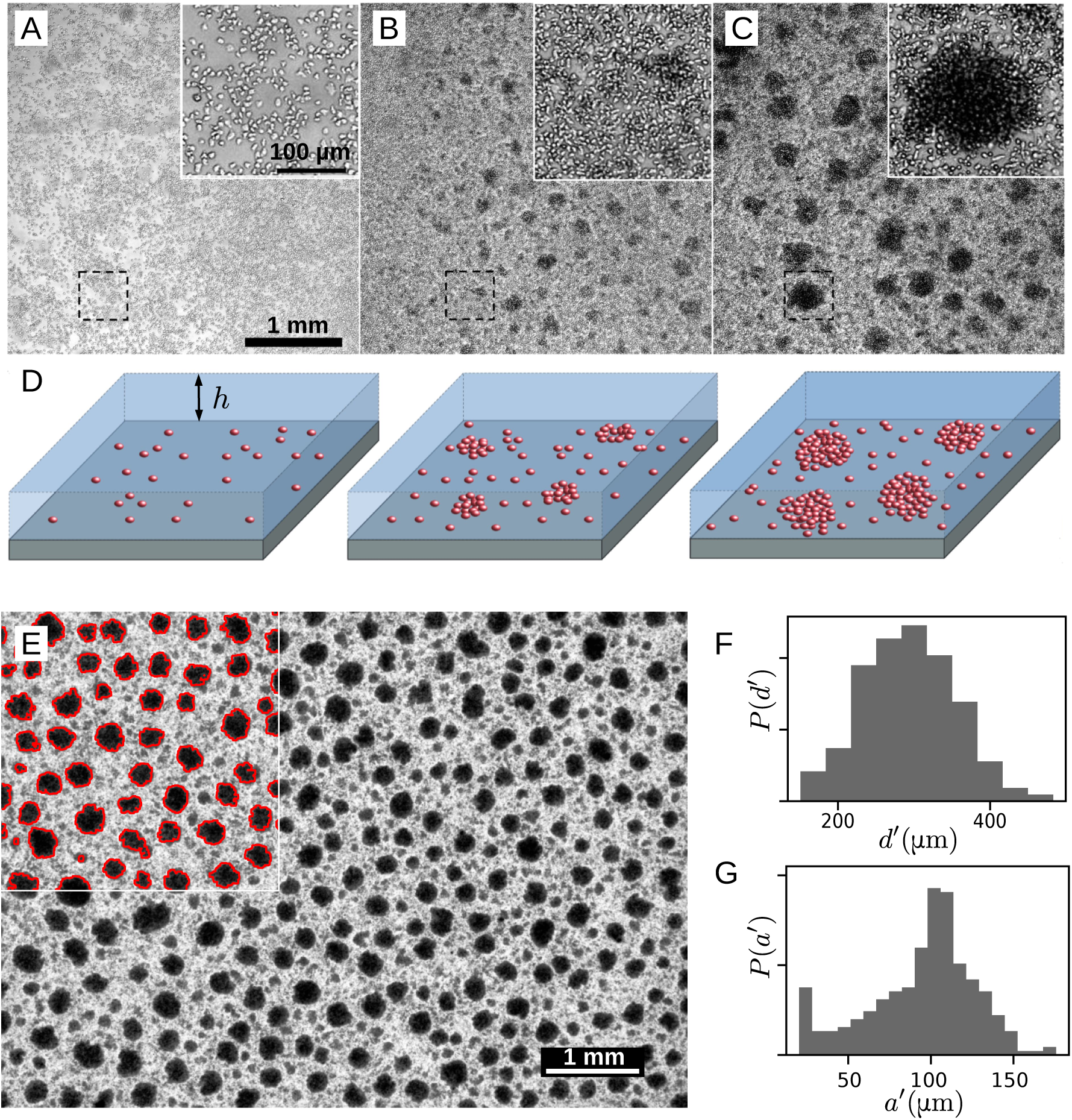
Microphase separation in a population of *Dictyostelium Discoideum* cells. (A-C) Brightfield images of the growth and aggregation process in the course of time. Each inset is a close-up of the dashed box. The liquid film height is *h* = 1.5 mm. (A) *t* = 24 h: the cell population has grown but remains rather homogeneous. (B) *t* = 48 h: while still dividing, cells gather in small growing aggregates. (C) *t* = 110 h: the aggregates have reached a steady state. (D) Sketch of the cell arrangement in the various stages. (E) Aggregates in steady state. Large field of view (8.2 × 6.1 mm^2^) brightfield image. Here *h* = 0.85 mm. The inset shows the contour of detected aggregates. (F-G) Probability distribution for the inter-aggregate distance *d′* and the aggregate radius *a′* computed from (E).

The striking observation is the self-organization of cells in compact, roughly circular domains with a finite length scale (Fig. 1E). Dark regions, indicative of packed, multilayered cell assemblies, exhibit both a typical size *a* and a typical center-to-center distance *d*. In Fig. 1F-G, one finds *a* = 100 ± 10 μm and *d* = 300 ± 60 μm and an aggregate thickness that usually remains below 40 μm (SM-C). The aggregates stand out on a light gray background of lower density with nearly confluent cells (Fig. 1C-D insets and movie M2). They are cohesive as they cannot be simply dissociated by gentle pipetting. If the calcium-dependent adhesion system is disabled, the domains dissociate (movie M3), proving that aggregates involve adhesion between cells. Remarkably, the self-organized state is not frozen (SM-B and movie M4): aggregates remain mobile over days and though their random motion may bring them in close vicinity, they rarely merge but rather avoid each other, suggesting that further growth is not limited by cell dynamics but prevented by an intrinsic repulsion.

The size of self-organized domains ranges from a few dozens to hundreds of micrometers and strongly depends on the liquid height above cells. A first hint is given by cell assembly below a meniscus (Fig. 2A). In the dish center, where the film is thinnest, an almost continuous domain is seen. However, as the liquid height increases upon approaching the wall, the giant domain is replaced by finite aggregates whose size decreases until complete disappearance. In subsequent experiments (Fig. 2B), cells were placed in distinct wells with different liquid height *h*. Here again, the thicker the liquid film, the smaller the aggregates.

**FIG. 2.**
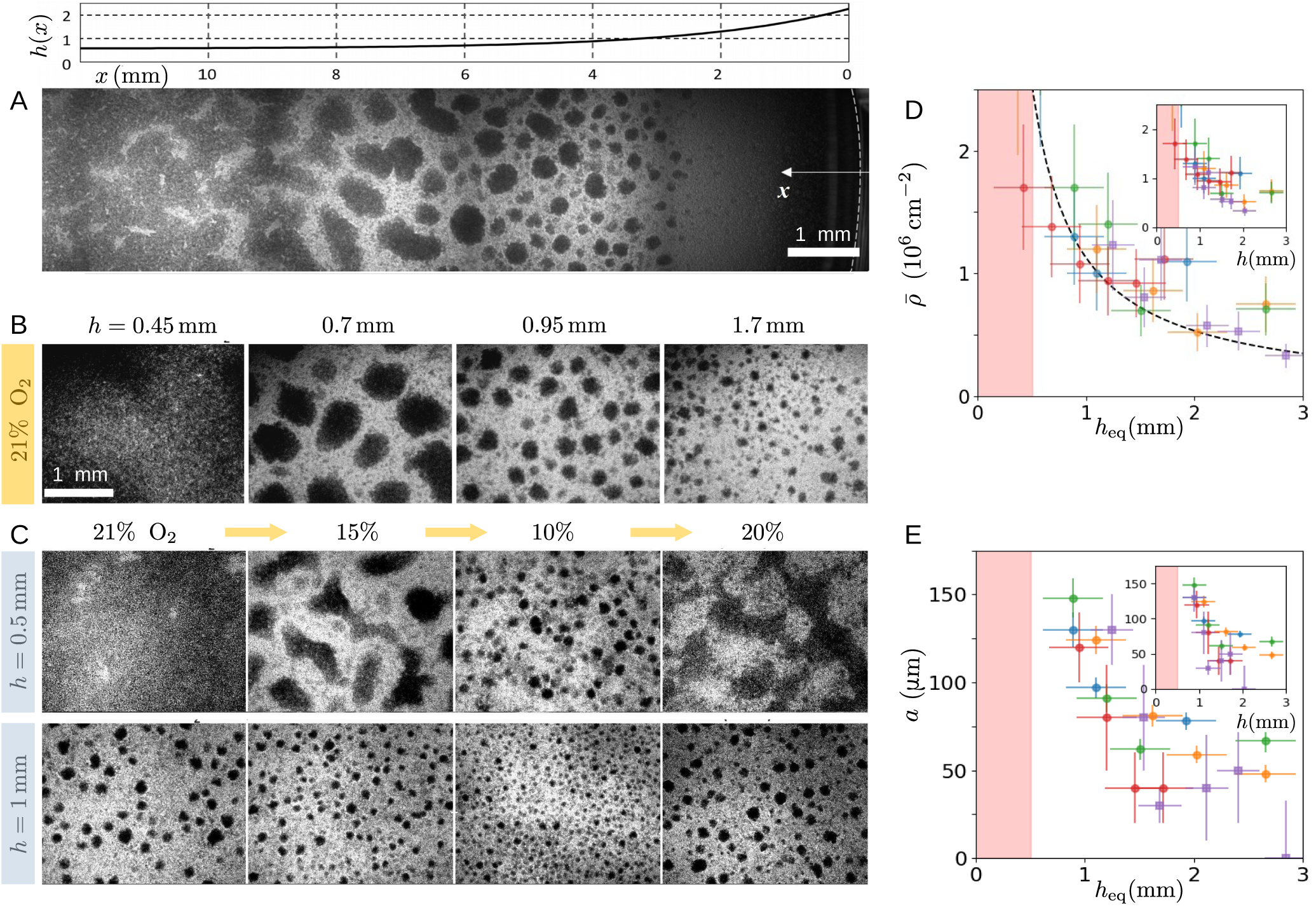
The size of aggregates is controlled by oxygen availability. (A) Mosaic image of the domains below the meniscus at the dish border. *x* is the distance from the dish wall and *h*(*x*) the height profile. (B) Influence of medium height *h* on the domain size in a normal atmosphere. (C) Effect of a step change in the oxygen content of atmosphere while keeping *h* fixed. Each step lasts at least 6.6 h. (D-E) Average (projected) cell density 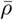 and typical aggregate radius *a* as a function of equivalent height 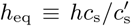 (main panels) or medium height *h* (insets). Colors indicate measurements from distincts samples. Circles and squares indicate oxygen partial pressure of 21% and 15% respectively. The dotted line in (D) is the prediction of Eq. (1). The pink area indicates the region where a single continuous aggregate forms.

Because the substrate is impermeable, the amount of oxygen available for cell consumption is governed by the film height *h*. Assuming a purely diffusive transport, the maximum flux of oxygen is *j*_m_(*h*) = *Dc*_s_*/h*, with *D* the diffusion coefficient and *c*_s_ the concentration at saturation which applies at the air-water interface. Using an atmosphere with controlled oxygen level (SM-A), we changed *c*_s_ to a value 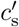 and observed that cell patterns were quite similar when the maximal flux *j*_m_ or the equivalent height 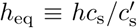 was the same, which suggests that the domain size depends primarily on the available flux of oxygen. To ascertain this point, aggregates having reached a steady state were submitted to a step change of atmosphere, a process equivalent to changing *h* but without inducing any disturbance on the cells or liquid above. The effect on aggregates is immediate: domains expand in enriched atmosphere (SM-E), and shrink in deprived atmosphere but recover their initial size upon going back to normal atmosphere (Fig. 2C and movie M5). Taken together, those observations demonstrate that oxygen availability, as quantified by *h*_eq_, is the key factor controlling the aggregate size.

To characterize our system, we obtain from direct measurements the medium saturation concentration *c*_s_ = 250 μM and the cell consumption rate *q* = 4.2 10^*−*17^ mol s^*−*1^ (SM-D). The average cell (projected) density 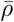 was found to depend on *h*_eq_: the thicker the film, the lower the number of cells (Fig. 2D). It is known that cell division is strongly suppressed in anoxic condition [26, 27]. Assuming that it stops entirely below a threshold *c*_div_, the density under a film of thickness *h* is

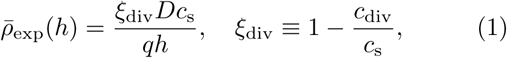

where the literature and our own measurements (SM-D) suggests *ξ*_div_ to be a few percent below unity. Taking *ξ*_div_ = 0.95 and using *D* = 2 10^*−*5^ cm^2^ s^*−*1^ [28], Eq. (1) yields 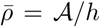, with 𝒜 = 1.05, 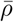 in 10^6^ cm^−2^ and *h* in mm, which is compatible with experimental data (Fig. 2D). The main observable characterizing the microphase separation is the typical aggregate size *a*, which depends on the equivalent height (Fig. 2E). For *h*_eq_ below 0.5 mm, the covering is continuous, corresponding to a formally infinite size (left side of Figs. 2A and 2B). When *h*_eq_ ≃ 1 mm, the largest aggregates that can be defined unambiguously are 150 μm in radius. For thicker films with *h*_eq_ = 2 − 3 mm, the aggregates are smaller, with radius around 50 μm. We now turn to the understanding of these experimental facts.

## A minimalistic model

We present a parsimonious approach capturing the main trends of experiments and involving only two basic assumptions: *(i)* cells spontaneously gather into aggregates with (projected) cell density *ρ*_a_, *(ii)* aggregates grow in size until the minimal oxygen concentration above them reaches a critical value *ĉ*. The former assumption is justified by cell-cell adhesion, a short-range attraction that promotes aggregation, whereas the latter acts a long-range repulsion, because oxygen depletion is stronger in larger aggregates. From this competition, a preferred domain size results. The microscopic mechanisms underlying such behavior need not be specified at this point.

Our model defines a pure diffusion problem, which in a simplified geometry of disk-like aggregates (Fig. 3A), can be solved analytically (SM-F). The minimal concentration is attained at the aggregate center (Fig. 3B-C) and reaches *ĉ* when

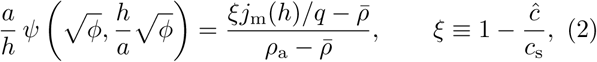

with *ϕ* the aggregate surface fraction and *ψ* a known function. As visible in Fig. 3D, the existence of aggregates is limited to a range of film thickness [*h*_min_, *h*_max_] with

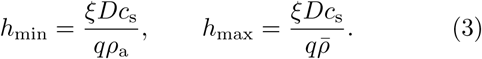

**FIG. 3.**
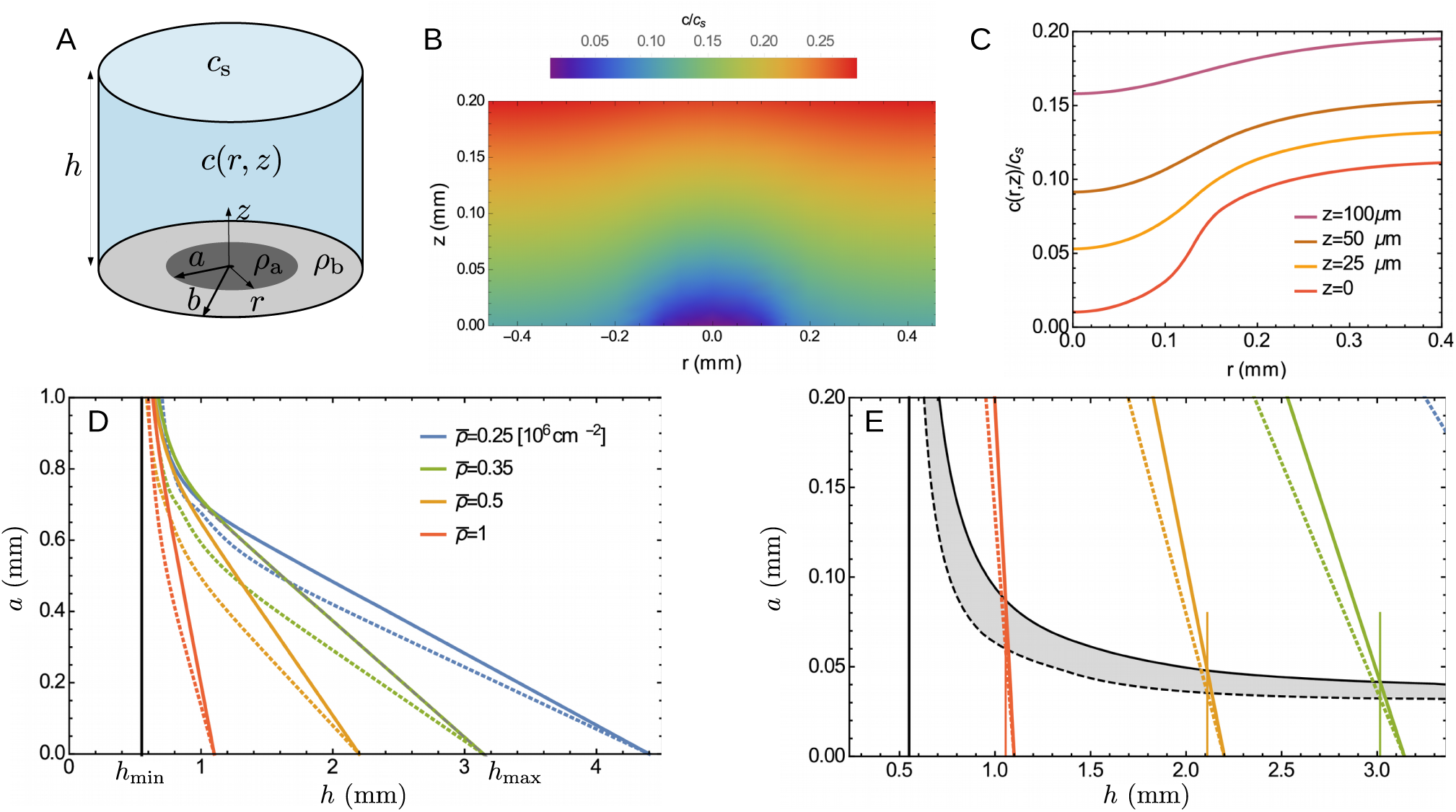
A simple model captures the domain size. (A) Geometry considered: The aggregate (resp. background) has radius *a* (resp. *b*) and cell density *ρ*_a_ (resp. *ρ*_b_). (B-C) Oxygen concentration field *c*(*r, z*) around an aggregate. The size domain *a* = 130 μm is the value such that the minimal concentration reaches the target concentration *ĉ*, that is *c*(0, 0)*/c*_s_ = *ĉ/c*_s_ = 0.01. Here 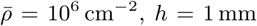 and *ϕ* = 0.4 *ϕ*_max_, where *ϕ*_max_ is the maximum aggregate surface fraction, reached when the background is empty of cells. (D) Aggregate size 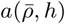 predicted when 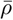 and *h* are independent parameters, as obtained from numerically solving Eq. (2). The continuous and dashed lines correspond to surface fraction *ϕ* = *ϕ*_max_ and 0.4 *ϕ*_max_ respectively. (E) The vertical lines indicate the film height *h* solution of 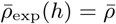 and the intersection point gives the aggregate size *a*(*h*) predicted when accounting for the thickness dependence of cell density. The black curves (with *ϕ* = *ϕ*_max_ and 0.4 *ϕ*_max_ for solid and dashed lines respectively) show the size expected in experiments.

At both ends of this interval, the system is homogeneous. For *h* = *h*_min_, the film is so thin that even with an infinite aggregate, the concentration *ĉ* is nowhere attained. For *h* = *h*_max_, even an aggregate of vanishing size makes the minimum concentration drop below *ĉ*. Near *h*_max_ where the aggregate size *a* is small, the solution of Eq. (2) can be expanded as *a* = *α*(*h*_max_ − *h*), with −*α* the slope and 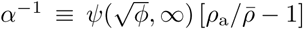. Such a linear approximation is actually excellent (Fig. 3D) and the intersection with the lower boundary *h* = *h*_min_ defines a characteristic aggregate size 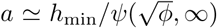, where 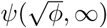 remains of order unity.

The predicted aggregate size is strongly dependent on the film height, especially at high mean cell density (Fig. 3D). With the parameters chosen (*ρ*_a_ = 2 10^6^ cm^−2^, *ĉ/c*_s_ = 0.01, see SM-F), the characteristic size *a* ≃ *h*_min_ is a fraction of millimeter, significantly above the experimental values. The reason is that 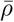 and *h* were treated as independent, whereas they are not. Accounting for the cell density 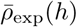 of Eq. (1) and introducing Δ*ξ* = *ξ* − *ξ*_div_ gives

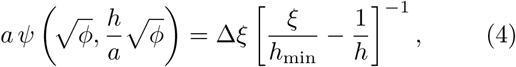

whose numerical solution is shown in Fig. 3E. In particular, for large *h/h*_min_, one finds *a ≃* Δ*ξ/ξ h*_min_ with Δ*ξ* = 0.04. The thickness dependence of cell density has two important implications. First, the typical aggregate size is not *h*_min_ any more but a significantly lower value around 50 μm, much closer to observations. Second, the aggregates can exist at large water heights. Both consequences originate from a single fact: the mean density spontaneously reached by the system corresponds to a liquid height slightly below the value *h*_max_ where aggregates would disappear (Fig. 3E).

Let us briefly consider the model predictions (Fig. 3D) for a step change in atmosphere when the cell density remains constant, since cells have no time to divide. When suddenly submitted to an oxygen-deprived atmosphere, aggregates should disappear. A clear reduction is observed (Fig. 2C and movie M5), though not a complete disappearance that we suspect is prevented by an increasingly slow dynamics. When submitted to an enriched atmosphere, aggregates are predicted to become larger, a trend that is unmistakably observed (SM-E) even if the final sizes are again difficult to ascertain. Thus, in spite of drastic approximations, our elementary model recapitulates the main features of aggregate behavior and supports the notion that oxygen regulation governs domain formation.

## Cell-based simulations

We now propose a microscopic description of the domain formation and illustrate how the picture of the analytic model can originate in the individual behavior of the cells. In our two-dimensional lattice model (Fig. 4A), a site can accommodate several cells, each of which obey three rules. First, cell-cell adhesion favors contact with their neighbors. Second, cells consume oxygen. Third, they exhibit aerotaxis toward oxygen-rich regions, but only in near anoxic condition when *c/c*_s_ *< c*_aer_ = 0.1, as demonstrated recently [29]. In our simulations, the oxygen concentration field, assumed steady, is evaluated with a Green function and the cells move according to a Monte Carlo (MC) algorithm with local moves only. A realization typically involves 10^4^ cells and 10^6^ − 10^7^ MC steps. The model definition, numerical method and parameters are detailed in SM-G.

**FIG. 4.**
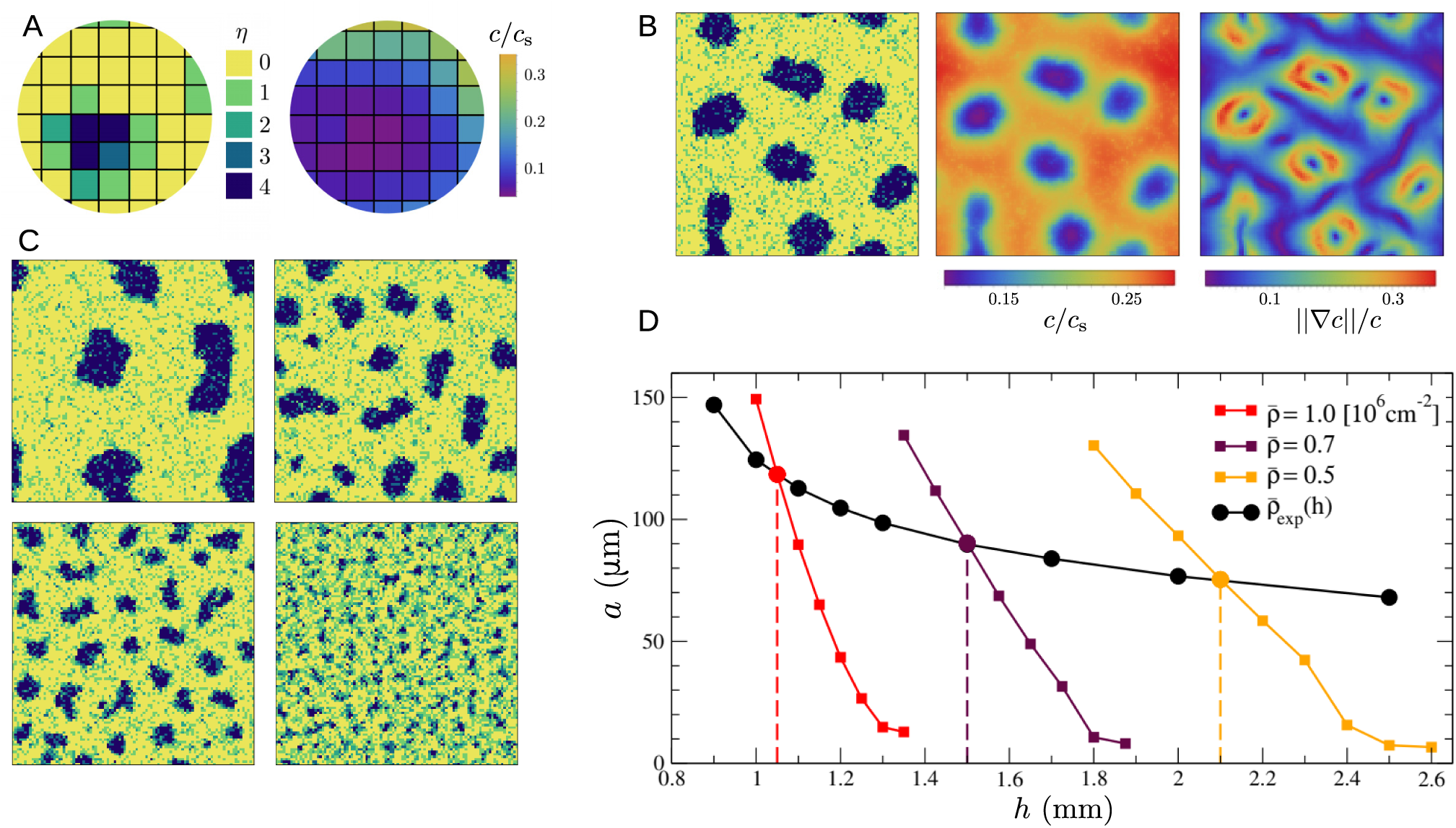
Cell-based simulations provide a microscopic view of microphase separation. (A) Schematic of the two-dimensional lattice model. Each site is occupied by a number *η* of cells, with maximum *η*_max_ = 4. The oxygen concentration field is assumed steady. Cells adhere to their neighbors, consume oxygen and become aerotactic at low concentration when *c/c*_s_ *< c*_aer_ = 0.1. (B) Example of aggregate. From left to right, occupation number, oxygen concentration and term ||∇*c*|| */c* controlling aerotaxis. Parameters: cell density 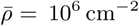, height *h* = 1.1 mm and lattice size *L* = 100. (C) The aggregate size depends on film height: simulations with *h* = 1.05, 1.15, 1.2, 1.3 mm yield an aggregate size *a* = 118, 65, 43, 15 ± 5 μm respectively. Here 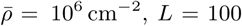. (D) Typical aggregate size *a* as a function of film height *h*, for several fixed cell density 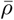 (colored squares). The vertical lines indicate the film height solution of 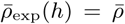. Also shown is the typical aggregate size expected in experiment (black dots). Lines are guides to the eye.

The competition between adhesion and aerotaxis is sufficient to drive the formation of finite-size aggregates (Fig. 4B-C). In the absence of aerotaxis, cells form aggregates, which further grow by fusion events. Such a coarsening process would lead, albeit slowly, to a divergence of domain size. In the presence of aerotaxis, the aggregate size converges to a finite value, indicating a stationary state. The microscopic mechanism can be inferred from Fig. 4B. Cell consumption induces oxygen depletion near aggregates, generating gradients that below *c*_aer_ trigger aerotactic escape and limit further domain growth. Consistent with this picture, the domain size is controlled by film thickness *h* and mean cell density 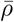, as visible in Fig. 4D. A comparison with Fig. 3D reveals that qualitatively the trends in the model and simulations are identical. Aggregates appear only below a height *h*_max_ and upon decreasing *h* from this point, their size increases linearly, with a slope controlled by the density 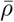. The simulations thus provide a microscopic basis for the behavior postulated in the minimalistic model.

For a quantitative comparison between simulations and experiments, we fix the cell size to 10 μm and take all other parameters from their known value, except the cell consumption *q*, which is reduced by a factor 1.2 (SM-G). Accounting for the height dependence of mean cell density, the aggregate size found in simulations is shown in Fig. 4D. As the film height varies from 1 to 2.5 mm, the aggregate radius decreases from 120 to 70 μm. This trend and the typical size around 100 μm are both consistent with observation data. Besides, as in experiments, the simulated aggregates never reach a static configuration but continually deform, move and rearrange (SM-G and movie M6). Such a feature is typically absent from inert systems and might be specific to microphase separation of living cells. All together, our cell-based simulations can capture the key features of aggregate behavior and confirm that oxygen-controlled domain formation can result solely from the interplay between adhesion, consumption and aerotaxis.

Quasi two-dimensional assemblies of *Dd* cells under liquid films can self-organize into domains of finite size because the combined effect of oxygen depletion and aerotaxis leads to an effective long-range repulsion that counteracts spontaneous aggregation. The mechanism is similar to the microphase separation of inert systems. There is, however, an interesting difference. Unlike purely twodimensional systems [19, 30, 31], the repulsion is mediated by the liquid film, whose equivalent thickness *h*_eq_ determines the range of effective interaction. Because *h*_eq_ is easily varied experimentally, it offers some control on the domain size, which can be changed dynamically. This is in stark contrast with block copolymers where the longrange repulsion is primarily governed by chain properties and is thus fixed once and for all [13].

Since domain formation in living cells has been observed in the past, it is worth pointing out how the involved phenomena, though superficially resembling, are actually different from the microphase separation reported here. The bacterial chemotactic patterns of Ref. [32] exhibit finite-size spots but in contrast with our aggregates, they are irreversibly “frozen in” and made of immobile bacteria. In the microcolonies of *Neisseria gonorrhoeae* [33, 34], where coarsening would eventually lead to a giant monodomain, the finite size is a consequence of kinetic slowdown and not the result of competing interactions. The stripe patterns of bacteria with density-dependent motility [35] involves a continuously expanding population, not a fixed number of cells. Finally, the classical aggregation of *Dd* results in a single millimetric immobile mound [36], clearly distinct from our smaller and dynamic aggregates.

The microphase separation of living cells opens questions and perspectives. First, we have focused on the domain size in steady state but many aspects call for in-depth exploration: the kinetics of domain formation and the interplay with cell division, the tridimensional structure of aggregates and their random motion, mode of migration and chemotactic ability [37, 38]. Second, an understanding of the biological mechanism underlying cell behaviors is needed. Third, we anticipate that the microphase separation demonstrated with *Dd* cells has a wider relevance because adhesion and aerotaxis are both widespread throughout the living world, from bacteria to higher eukaryotic cells [39, 40].

We conclude on the broader significance of our findings in the context of biological patterning and morphogenesis. Embryonic development was long attributed to an inductive process governed by exogenous signals such as maternal chemical gradients or morphogens [41]. In contrast, a recent line of work with embryoid models emphasizes the evidence for endogenous self-organization [42]. The two mechanisms may actually act concurrently, a combination termed guided self-organization. The oxygen-controlled microphase separation found here offers a striking instance of such a hybrid process. No pattern is imposed from outside but the availability of oxygen, that may be controlled externally, dictates whether cells aggregate or not and which domain size is ultimately selected. Interestingly, a related approach of “directed assembly” has been exploited for block copolymers [43]. Guided self-organization is a thus generic mechanism relevant both in technological applications and in the natural realm. Finally, multicellular aerobic life is inevitably constrained by the fact that diffusion is slow over large distances [44, 45]. While the genesis of multicellularity remains debated [46], one may wonder whether microphase separation is involved and possibly reassess the role of oxygen in pattern formation and life evolution. We expect physical concepts to be instrumental in this endeavour.

This research was funded by grants to J-P. R., from CNRS (MITI, Défi Modélisation du vivant, 2019), from ANR (ADHeC project, ANR-19-CE45-0002-02, 2019), and from International Human Frontier Science Program Organization (RGP0051, 2021).

## Supplementary Material

**SM-A**: Materials and Methods

**SM-B**: Steady state

**SM-C**: Aggregate structure

**SM-D**: Oxygen measurements, cell consumption and division

**SM-E**: Aggregate response to a step change in oxygen

**SM-F**: Analytical model

**SM-G**: Simulations of a cell-based model

**SM-H**: Movies

## A. MATERIALS AND METHODS

### 1. Cell preparation and live cell imaging

AX2 axenic cells of *Dictyostelium discoideum* were grown in HL5 medium with glucose (Formedium, Norfolk, UK) at 22°C in tubes with shaking conditions (180 rpm for oxygenation). Exponentially growing cells were harvested, counted and deposited in 6-well plates, with a initial surface cell density *ρ*_0_ between 5 10^3^ and 2 10^5^ cells/cm^2^. Cells were submerged below liquid growth medium with height ranging from *h* = 0.5 to 3 mm. The temperature was kept constant at 22°C.

The growth and aggregation of *Dd* cells were observed in transmission with three types of microscope: *(i)* a TE2000-E inverted microscope (Nikon) equipped with a motorized stage, a 4X Plan Fluor objective lens (Nikon) and a Zyla camera (Andor) using brightfield for most of the timelapse experiments lasting up to several days (Figs. 1A-C, 2B-C, S6, and movies M1, M3 and M5), *(ii)* a binocular MZ16 (Leica) equipped with a TL3000 Ergo transmitted light base (Leica) operated in the one-sided darkfield illumination mode and a Leica LC/DMC camera (Leica) for large field experiments (Figs. 1E, S1C, S2 and movie M4) and finally *(iii)* a confocal microscope (Leica SP5) with a 10X objective lens for larger magnification experiments (Fig. S3A-C and movie M2) and to create large field reconstituted mosaic images (Fig. 2A).

### 2. Atmosphere with controlled oxygen level

In several experiments (Figs. 2B-C, Fig. S6), we varied the oxygen concentration in the atmosphere surrounding the liquid film. Plates were placed in a home-made environmental chamber fitting our microscope stage with a pair of top and bottom glass windows. A mixture with pure N_2_ (0% O_2_) and air (21% O_2_) was prepared in a gas mixer (Oko-lab 2GF-MIXER, Pozzuoli, Italia) and injected continuously in the chambers at about 300 mL/min, so that the oxygen level in the atmosphere can be maintained at a prescribed value.

### 3. Medium height profile

The medium height *h* in dish center was estimated as a function of time by dividing the corrected volume by the dish area, typically 9.6 cm^2^ in 6-well plates. The corrected volume is the initial culture medium volume from which are subtracted the evaporated volume *V*_evap_ and the meniscus volume *V*_menis_. The evaporated volume *V*_evap_ was deduced by measuring the remaining sample volume in a precise serological pipet at the end of each experiment. It was extrapolated to any time assuming a constant evaporation rate. From the height profile in the small slope approximation *h*(*x*) = *h* + *δh*(0) exp(−*x/L*_c_), the meniscus volume was computed as *V*_menis_ = 𝒫*δh*(0). Here 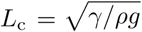 is the capillary length [A1], 𝒫 is the dish perimeter, 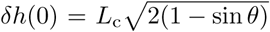 is the meniscus height at the dish wall (*x* = 0) relative to the medium height *h* in the center and*θ* is the contact angle. The same small slope approximation was used to plot the meniscus profile of Fig. 2A. The surface tension, measured with a Langmuir balance (NIMA, England), is *γ* = 55 mN/m. The contact angle value *θ* = 48 ° was obtained for a 3-day sample and used for the remainder of the study. Its value was periodically verified by taking side view images of the meniscus.

### 4. Image analysis

#### Aggregates

A home-made algorithm in Python was used to detect the domains. A typical image is shown in Fig. 1E of the main text, where the boundary of identified aggregates is drawn in red. We note that some clumps of cells, with small or moderate size and characterized by intermediate gray level, may or may not be identified as aggregates. The domains with large size, which are most often very dark, are detected unambiguously, i.e. they are not sensitive to the precise parameters used in the algorithm. They thus define a robust population of domains, which is the focus of our study.

Once aggregates are detected, we compute their area 𝒜, their equivalent radius 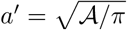 and identify the typical radius *a* from the maximum in the distribution *P* (*a′*). We choose the maximum and not the mean of the distribution, because the former is independent of values found for small aggregates whereas the latter is not. Finally, we also compute for each aggregate the distance *d* to the nearest neighbor and use the mean *d ′* of the distribution *P* (*d′*) as a typical inter-aggregate distance.

### 5. Cell density

#### Average cell density

A the end of each experiment, cells were detached by thorough pipetting, the cell suspension in liquid medium was immediately collected with a serological pipet and part of it was used in a hemocytometer to obtain the average (projected) cell density 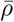. The counting was repeated twice with two different cell suspension sample volumes to estimate the counting error.

#### Background cell density

The background is defined as the area outside identified aggregates. The cell density *ρ*_b_ in the background can be determined by automatic detection of individual cells on images using the Find-Maxima plugin in ImageJ.

## B. STEADY STATE

To ascertain that our system has reached a steady state, we considered several quantities: the background phase cell density, the mean aggregate size and the distribution of aggregate size.

Figure S1A shows a semilog plot of the background cell density *ρ*_b_ as a function of time for the same medium height *h* = 1.5 mm but starting from different initial cell densities *ρ*_0_. Among the three independent experiments, the red curve corresponds to the experiment displayed in Fig. 1A-C of the main text (*ρ*_0_ = 7.5 10^4^ cm^−2^). The doubling time in the exponential growth phase is similar for the three experiments, with a value 8 ± 1 h. The background cell density after aggregation (indicated by arrows) is constant at about 5 ± 1 10^5^ cm^−2^.

The size of aggregate (Fig. S1B) also exhibits a plateau reached about 24 h to 36 h after the onset of aggregation. The red curve corresponds again to the experiment of Fig. 1A-C (*h* = 1.5 mm). The blue curve corresponds to the experiment of Fig. 1D-F of the main text (*h* = 0.85 mm, *ρ*_0_ = 10^5^ cm^−2^). For the latter, we show in Fig. S1C two snapshots of the aggregates at times separated by 10 h. To the naked eye, there is no obvious change in the domain size. This can be confirmed quantitatively by computing the distributions of size and inter-aggregate distance (Fig. S1D-E). In both cases, there is no change in the histogram bin where the maximum is reached and the increase of the mean average size is only 5% after 10 h. To a very good approximation, one can consider the aggregate have indeed reached a steady state.

**FIG. S1.**
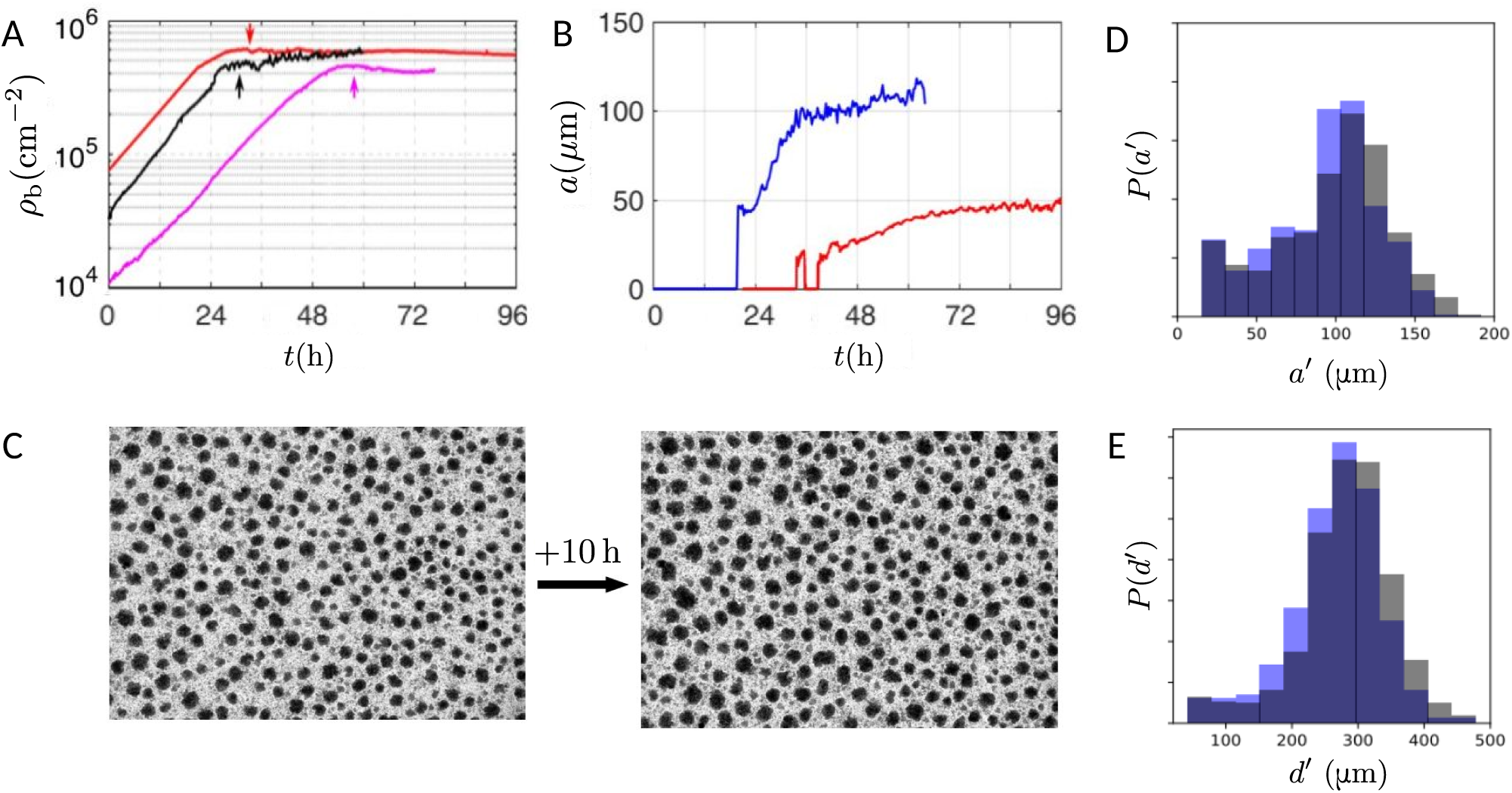
(A) Time evolution of the background cell density *ρ*_b_ The medium height is *h* = 1.5 mm and the initial cell densities are *ρ*_0_ = 1.2 10^4^, 2.2 10^4^ and 7.5 10^4^ cm^−2^ for pink, black and red curves respectively. (B) Time evolution of the mean aggregate size *a*. The medium height is *h* = 0.85 and 1.5 mm for blue and red curves respectively. (C) Snapshots of the experiment corresponding to the blue curves in (B) after *t* = 53 h and 10 h later. (D-E) Superimposed distributions of the domain size *P* (*a′*) and first neighbor’s distance *P* (*d′*) for the two snapshots at *t* = 53 h (blue) and 10 h later (gray).

Remarkably, aggregates themselves remains very mobile and never settle into a static configuration. This is illustrated in movie M4 and in Fig. S2, which shows the trajectory of many aggregates over one day. Aggregates actually move over a distance much larger than the typical inter-aggregate distance, and do so without coalescing or merging with their neighbours.

**FIG. S2.**
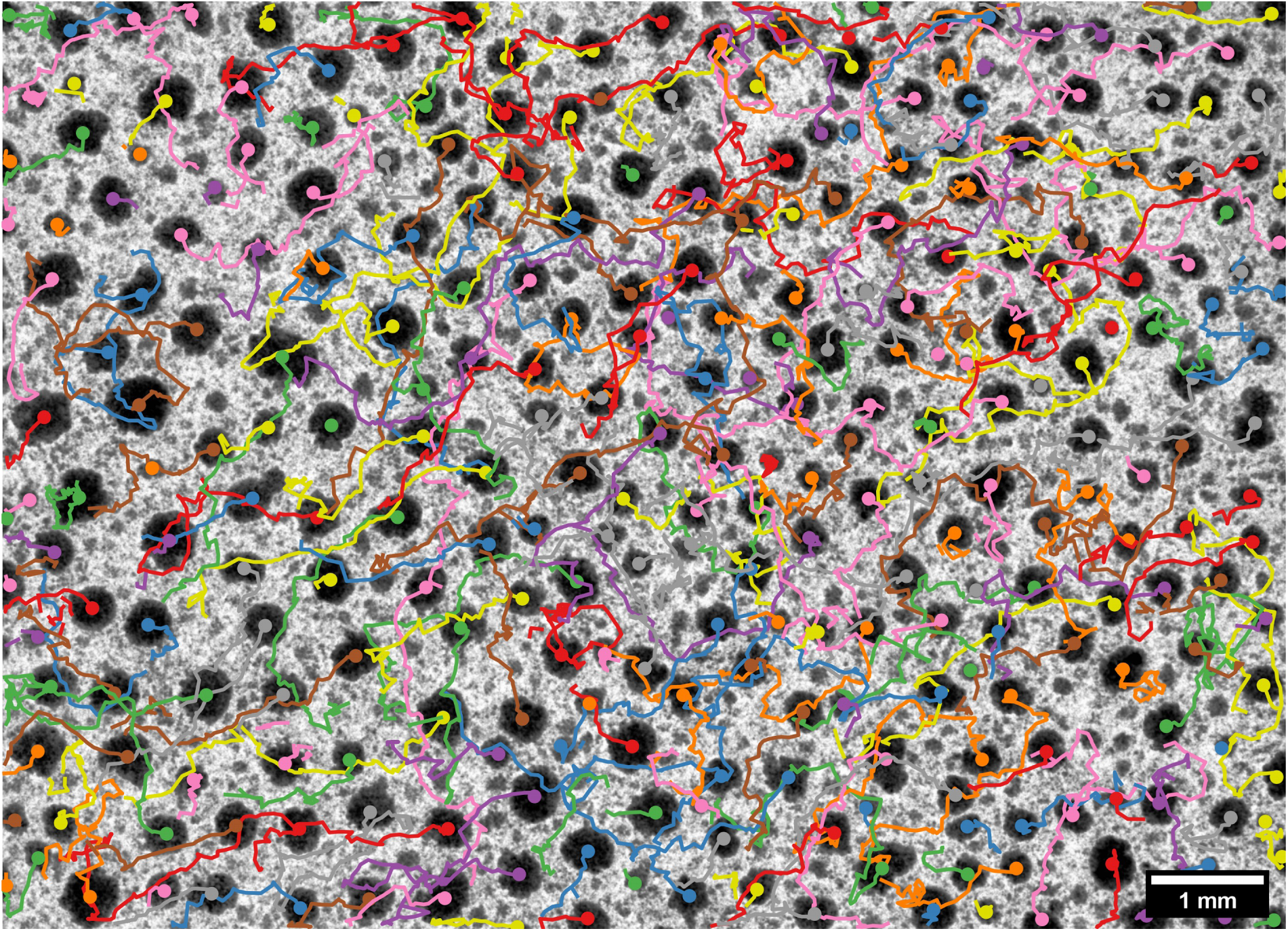
Trajectories of domains tracked over 24 h. Each dot indicates the final position. The film height is *h* = 0.85 mm, as in Fig. 1E.

## C. AGGREGATE STRUCTURE

### 1. Aggregate cohesion

*Dd* cells in development stage in starvation buffers are known to possess three adhesive systems [C1]. The first to be expressed is calcium dependent and can be disabled by EDTA (Ethylenediaminetetraacetic acid), a chelator of divalent ions.

Once aggregates reached their steady state, we added 20 μL EDTA at 500 μM in one well of a 6-well plate with about 2 mL of medium. The injection was made drop by drop as gently as possible to avoid any disturbance. The aggregates then start dissociating (movie M4), a process seen immediately in the injection zone and elsewhere in the subsequent hours. These observations indicate that the calcium dependent adhesive system is operative in a nutrient buffer at high cell density and that cell-cell adhesion is responsible for the aggregate cohesion.

### 2. Aggregate height

We first examine a typical aggregate, whose radius 70 μm is close to the typical value *a* = 65 ± 25 μm obtained with medium height *h* = 1.5 mm. To visualize the three-dimensional structure, we used Z-stacks in transmission mode using 10X objective lens on a confocal (Leica SP5, Germany). The reference slice at *z* = 0.00 μm (Fig. S3A) shows the bottom of the aggregate surrounded by a first layer of cells spread on the substrate (yellow arrows) and not very mobile (movie M2). At *z* = 7.11 μm above the reference (Fig. S3B), a second layer of cells, more rounded and very mobile is present (red arrows). At *z* = 30.81 μm (Fig. S3C), we detect the aggregate upper boundary. Taking into account the uncertainty in the manual estimation of reference level (±2 μm) and of the upper boundary (±7 μm), this suggests a height around 30 ± 7 μm.

A second indication of aggregate height is provided by aggregates climbing PDMS vertical pillars, as illustrated in Fig. S3D. Most aggregates had a height within 30 μm but aggregates as thick as 40 μm were occasionally encountered.

**FIG. S3.**
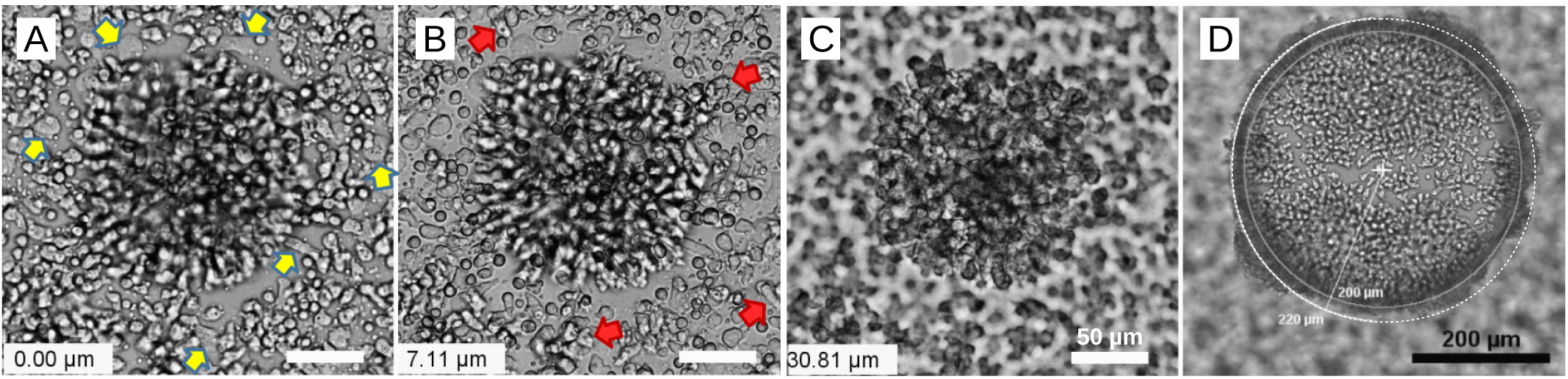
Multilayered structure of a typical aggregate (70 μm in radius, medium height *h* = 1.5 mm). (A-C) Confocal slices at (A) 0.00 μm, (B) +7.11 μm and (C) +30.81 μm. Yellow arrows indicate some cells of the first layer (not mobile) and red arrows indicate cells of the second layer (rounded and mobile). (D) Side view of aggregates climbing PDMS vertical pillars. The pillar is 200 μm in radius, as shown by the inner circle. The outer circle plotted is at distance 20 μm from the inner circle. The dark layer surrounding the inner circle gives a side view of aggregates.

### 3. Aggregate cell density

To estimate the projected cell density in aggregates *ρ*_a_, we use the relation 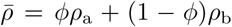. We consider the typical system shown in Fig. 1A-C and Fig. 2D with *h* = 1.5 mm of medium. From the measured mean cell density 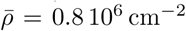 the background cell density *ρ*_b_ = 0.5 10^6^ cm^−2^and the surface fraction *ϕ* = 0.2, one finds *ρ*_a_ ≃ 2 10^6^ cm^−2^. Because of large error bars, the actual value may actually fall in large range around this average, but this provides at least a reasonable estimate.

## D. OXYGEN MEASUREMENTS, CELL CONSUMPTION AND DIVISION

### 1. Oxygen measurements

Oxygen concentration were obtained using a commercial optical “robust oxygen probe” (OXROB3) coupled to its oxymeter (Firesting, Pyroscience, Aachen, Germany). The measurement is based on the luminescence quenching by oxygen of a redflash indicator deposited at the probe tip.

We systematically controlled the oxygen content in our homemade environmental chamber. More specifically, we measured the gaseous volume percentage, defined as 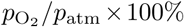, where 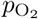 and *p*_atm_ are respectively the partial pressure of oxygen and the barometric pressure of ambient air.

To obtain the oxygen concentration at saturation in the HL5 *Dd* growth medium under normal atmosphere, the probe was plunged in 6-mL bottle filled with the medium but no cells. At temperature 22°C, we found *c*_s_ = 250 ± 20 μM, in agreement with literature values [28].

### 2. Consumption of individual cells

To measure the oxygen consumption of cells, we filled a 6-mL glass bottle with a concentrated cell suspension, which contains typically 2 10^7^ cells, and closed it with a cap through which the oxygen probe was plunged in the liquid. The cap was carefully sealed around the probe to avoid external oxygen entering the bottle. During the time required to prepared the cell suspension and the oxygen probe, the oxygen concentration inside the bottle dropped to about 200 μM. Then it linearly decreased to nearly zero in about 1600 s, as shown in Fig. S4A. The cell oxygen consumption which is proportional to the slope of this curve is nearly constant (Fig. S4B), with a value of *q* = 4.2 10^*−*17^ mol s^*−*1^ cell^*−*1^ (average over six experiments), in a concentration range extending from 150 μM down to a few μMs. For lower concentration, the consumption drops abruptly. The typical concentration at which cell consumption becomes concentration dependent is thus in the range 2.5 − 10 μM, which corresponds to *c*_csm_*/c*_s_ in the range [0.01, 0.04].

**FIG. S4.**
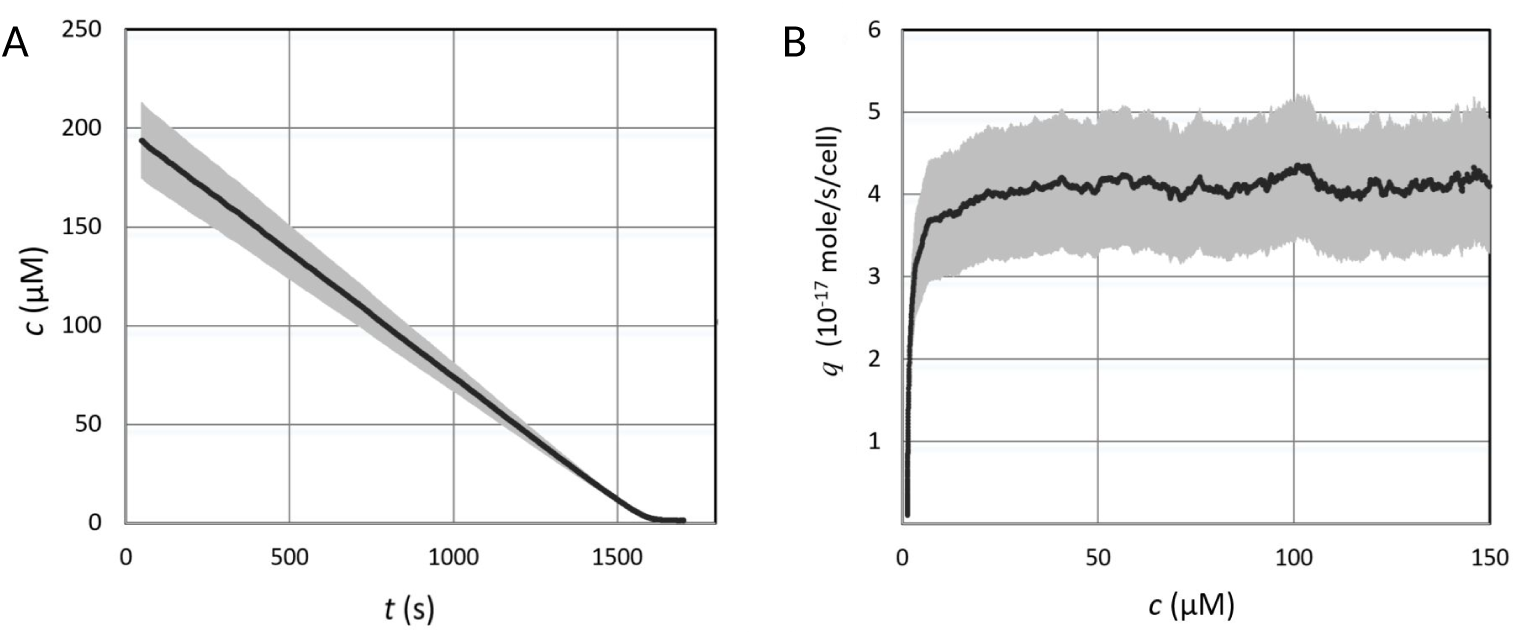
Measurement of oxygen consumption by AX2 cells. (A) Change in oxygen concentration as a function of time due to cell consumption. (B) Deduced cell consumption as a function of the oxygen concentration. The black solid line represents a typical experimental measurement. The shaded grey region indicates the standard deviation over six measurements (*N* = 6).

### 3. Cell division at low oxygen concentration

Starting from very low cell densities at about 3000 cm^−2^, we monitored the cell growth by timelapse microscopy under various atmosphere oxygen levels in our environmental chamber. Given the low cell densities which make total consumption small, the dissolved oxygen concentration in the culture medium is simply fixed by the oxygen level outside. Figure S5A shows a typical growth at two very different oxygen levels. In the initial atmospheric reference condition (21% O_2_, *c* = *c*_s_ = 250 μM), the growth is exponential with division time *T*_div_ ≃ 8 h (i.e. 4-fold increase during 16 h), in agreement with previous measurements [D1]. From 22 h to 45 h, pure N_2_ was injected in our chamber up, leading to a residual level of 0.15% O_2_ (*c* = 1.8 μM). During the first 6 hours of this severe hypoxic condition, the population continues to grow although at a decreasing rate. The cell number then reaches a plateau and eventually slightly decreases [D2]. Upon re-injection of 21% O_2_ air at time 45 h, the population still slightly decreases during about 6 h, but it later recovers and grows again. These observations indicate that at *c* = 1.8 μM, cells finish their life cycle before entering a resting phase (quiescence). The existence of the 6 hours lag period upon air re-injection implies that after quiescence, cells need time to resume their normal life cycle.

We tested a range of oxygen levels and observed that the growth index, that is the fold change in the number of cells between between *t* = 0 h and *t* = 16 h, is changing with oxygen concentration (Fig. S5B). In particular, at *c* = 5 μM and *c* = 12.5 μM, the growth index is *N* (16 h)*/N* (0 h) = 1.2 and 2 respectively. Our values (squares) are in agreement with literature values [26, 27] for the AX3 cell lines (triangles and circles respectively). Interpolating the data with a simple exponential form (dotted line) leads to the estimate *c*_div_ ≃ 10 μM for the concentration at which the growth is divided by two with respect to the reference in normoxic conditions. Accordingly, we use *c*_div_*/c*_s_ = 0.05 in Eq. (1) of the main text.

**FIG. S5.**
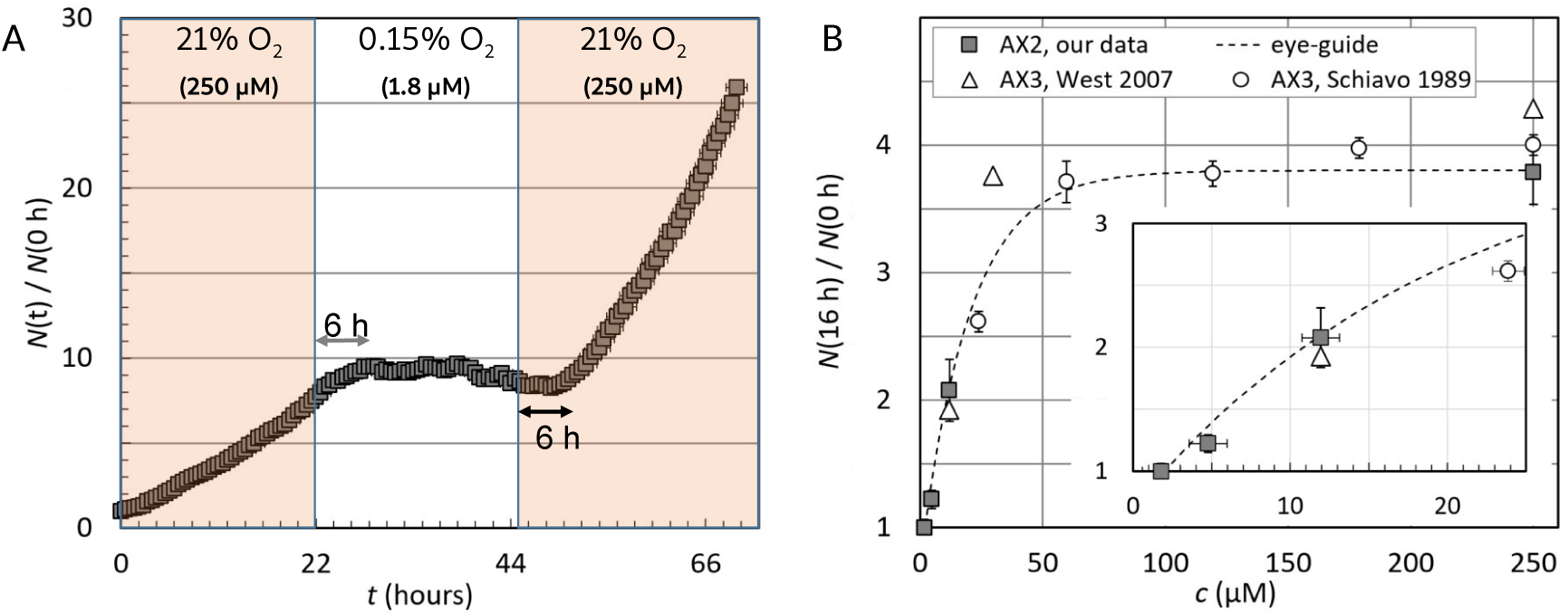
Oxygen dependence of cell growth. (A) Number of cells *N* (*t*) as a function of time, with changing oxygen levels. (B) Dependence of growth index on oxygen concentration in culture medium. The growth index is the fold change in the number of cells between *t* = 0 h and *t* = 16 h. Filled squares show our measurements with the AX2 cell line (number of measurement is 3, 4, 3 and 7 for *c* = 1.8, 4.8, 11.9 and 250 μM respectively). Triangles and circles correspond to literature value with AX3 cell lines [26, 27] interpolated from 24 h to 16 h assuming an exponential growth. The inset shows the 0 − 25 μM region. The dotted line is a guide for the eye.

## E. AGGREGATE RESPONSE TO A STEP CHANGE IN OXYGEN

We show in Fig. S6 the effect of increasing the amount of oxygen available to cells. Images are separated by 5 hours, a time interval during which cell division is negligible [E1]. Whatever the initial state fixed by the medium height *h* – large or small aggregates –, the typical size of domains substantially increases.

**FIG. S6.**
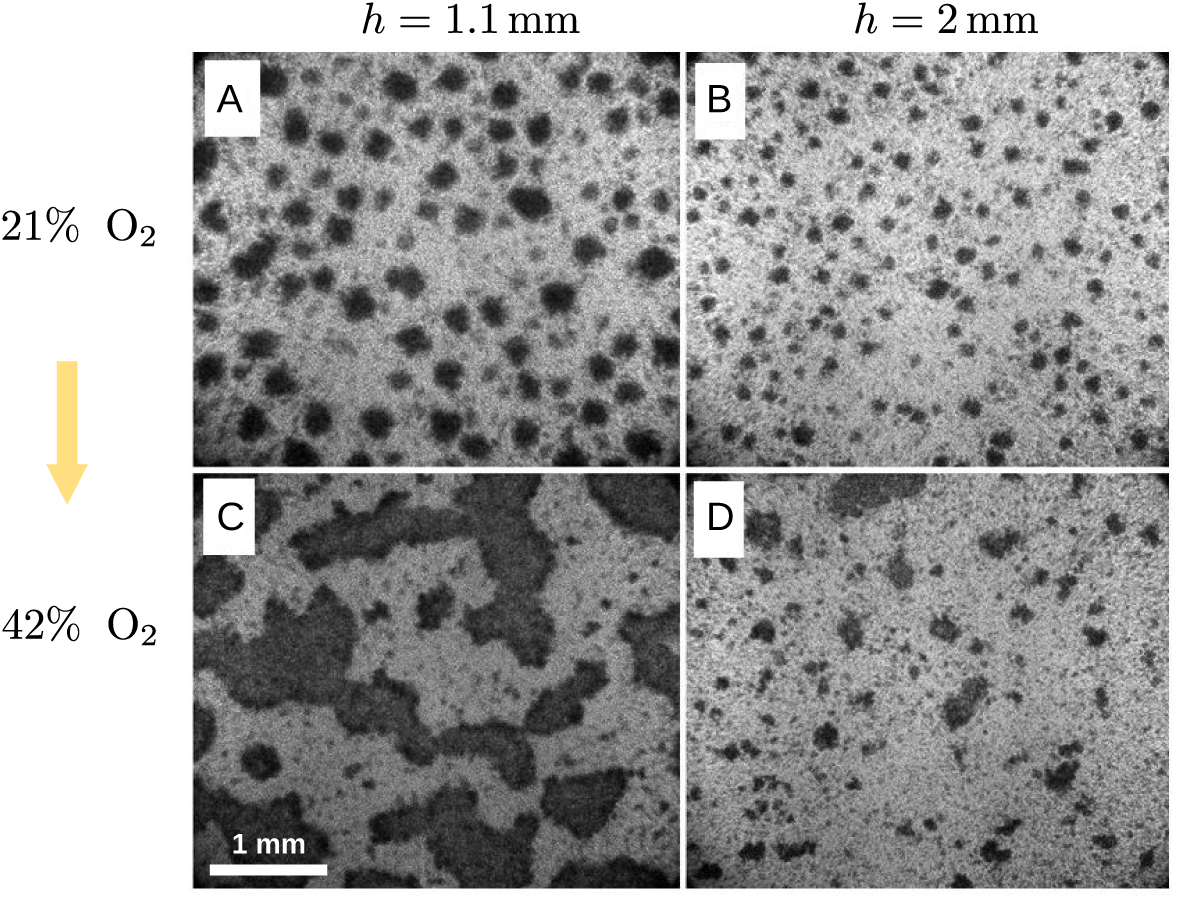
Effect of a step increase of oxygen on the aggregates. The oxygen level in the atmosphere above the liquid medium is doubled with respect to normal conditions. (A-B) Bright field images of aggregates formed under normal atmosphere and two medium heights. (C-D) Images taken 5 h after the step change.

## F. ANALYTICAL MODEL

### 1. Model definition

Our simplified model for aggregate is shown in Fig. S7. An aggregate is represented as a disk of zero thickness and radius *a* placed on the bottom surface. The cell density is *ρ*_a_ within the aggregate and *ρ*_b_ in the background outside. Because the mean cell density 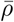 will be fixed in the following, an aggregate can grow in size only by depleting the surrounding region. The individual cell consumption *q* is assumed constant. The oxygen flux consumed by the cells is *J*_a_ = *qρ*_a_ and *J*_b_ = *qρ*_b_ on the aggregate and background respectively. The aggregate is not isolated but surrounded by identical neighbours arranged on a hexagonal array. To make the calculation analytically tractable, the hexagonal boundary of the unit cell is replaced by a disk of radius *b* of equal area, so that azimuthal invariance is recovered. A vanishing lateral flux is imposed at this effective boundary. The concentration on the top surface is fixed to the saturation value *c*_s_. The surface fraction of aggregate is *ϕ* = *a*^2^*/b*^2^ and the mean density 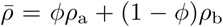. We will assume that the aggregate cell density *ρ*_a_ is a fixed value, independent of aggregate size and surface fraction.

**FIG. S7.**
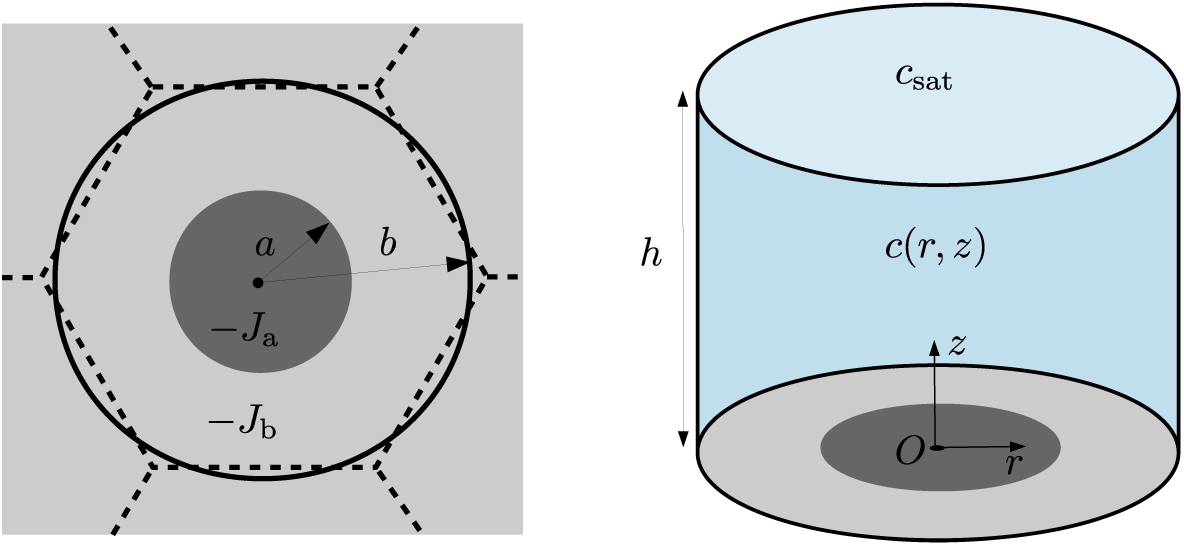
Configuration considered in the analytical model: an aggregate is represented as a disk of radius *a*, below a liquid film of height *h*. The flux of oxygen consumed is *J*_a_ on the aggregate and *J*_b_ on the background outside. We compute the concentration at the origin and seek for which aggregate radius *a* it reaches the target concentration *ĉ*.

### 2. Derivation of analytical solution

Here we give the details of the derivation for the oxygen concentration field *c*(*r, z*), which is a purely diffusive problem. The notations are shown in Fig. S7. It is convenient to work with the function

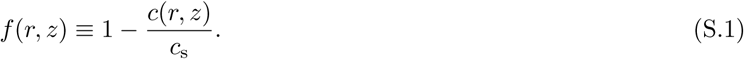

Unless otherwise mentioned, we chose units so that *h, D* and *c*_s_ are all unity. The equation and boundary conditions to be satisfied by *f* (*r, z*) are

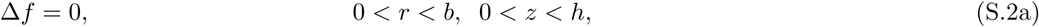

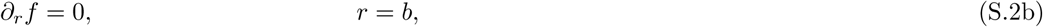

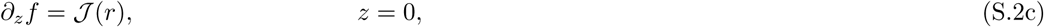

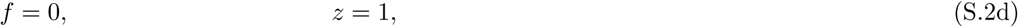

where 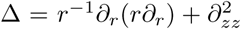 denotes the Laplacian. The flux perpendicular to the lateral wall is taken as zero and 𝒥 (*r*) is the arbitrary flux imposed on the bottom surface. Applying classical methods [F1], the solution is

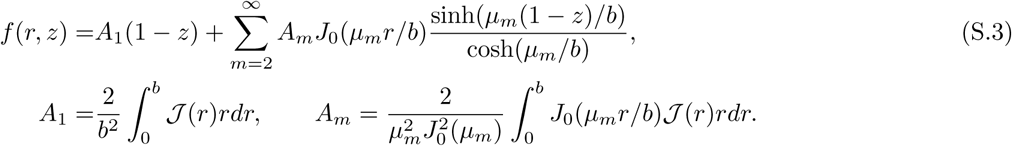

Here, *J*_*n*_ is the Bessel function of order *n, μ*_*m*_ is the *m*th zero of 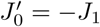, with the convention *μ*_1_ = 0. We consider an imposed flux

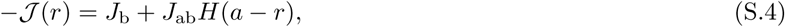

with *H* the Heaviside function, *J*_ab_ ≡ *J*_a_ − *J*_b_ and 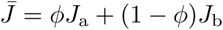 is the mean flux. We then find for the value at the bottom surface

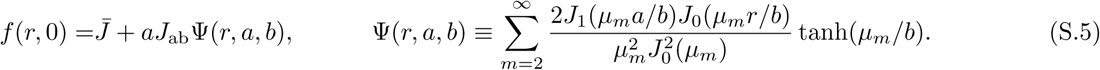

Now, putting back dimensions, the condition that the minimal concentration, obtained at *r* = 0 and *z* = 0, is the target value *ĉ* can be rewritten as

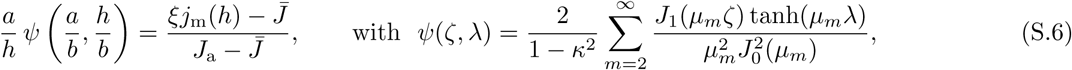

where we used 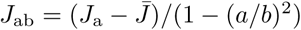. Since fluxes are proportional to cell densities, Equation (S.6) gives back Eq. (2), where 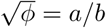 is assumed fixed.

The function *ψ*(*ζ, λ*), computed numerically, is plotted in Fig. S8. For *λ* = *h/b >* 1, the dependence on *λ* is negligible and *ψ*(*ζ, λ*) can be approximated as *ψ*(*ζ*, ∞). Taking *ζ* in the range [0.1, 1] includes all surface fraction of interest since *ϕ* = *ζ*^2^. In this domain, the function *ψ*(*ζ*, ∞) has limited variation, being confined in the interval [0.38 − 0.94] and remains a prefactor of order one. Note that this conclusion holds in most of the relevant parameter space but not in the vicinity of *h*_min_. In this region, the domain size *a* diverges while 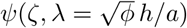 approaches zero as *λ* → 0.

**FIG. S8.**
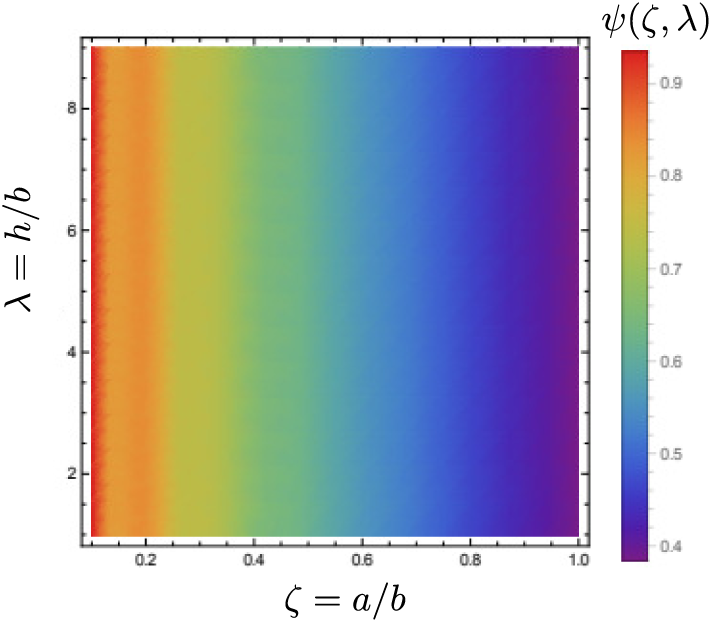
Function *ψ*(*ζ, λ*) defined in Eq. (S.6).

### 3. Choice of parameters

Here we explain the parameters taken in applying the model. The values for *c*_s_, *D* and *q* were introduced in the main text. The saturation concentration *c*_s_ = 250 μM and the cell consumption rate *q* = 4.2 10^*−*17^ mol s^*−*1^, were both measured experimentally as detailed in SM-D, The diffusion coefficient is *D* = 2 10^*−*5^ cm^2^ s^*−*1^ [28]. For the aggregate (projected) cell density, a reasonable estimate is *ρ*_a_ = 2 10^6^ cm^−2^ (SM-C) which corresponds to *h*_min_ = 0.55 mm according to Eq. (3). As regards the critical concentration *ĉ*, the steady state we consider has constant cell number because cells have stopped dividing (Fig. S1A), which suggests a concentration below *c*_div_ everywhere, and a significantly lower value at the aggregate center where it is minimal. We therefore fix *ĉ/c*_s_ = 0.01 and note that this value could be doubled or halved with comparable results. Finally, the surface fraction *ϕ* remains a free parameter. It is bounded by the maximal value *ϕ*_max_ reached when all cells are inside aggregates and none outside (empty background with *ρ*_b_ = 0). It turns out that the choice of *ϕ* between *ϕ*_max_ and lower values (0.4 *ϕ*_max_ for instance in Fig. 3E) has only a limited influence on the results, as visible in Fig. 3E.

## G. SIMULATIONS OF A CELL-BASED MODEL

### 1. Model definition and parameters

We introduce a minimal model with two basic ingredients. The first is adhesion between cells that are in contact, which favors the formation of aggregates. The second is aerotaxis, which drives motion along the gradient of oxygen and opposes the growth of large domains.

The model is lattice-based and represents each cell individually. A cell occupies a single site of a two-dimensional square lattice. To account in an effective manner for the three-dimensional structure of the aggregates, which may have several layers, a site can accommodate several cells. The number *η*_*m*_ of cells at site *m* is limited to a maximum value *η*_max_, which reflects the limited height of aggregates. We define the neighbors of a site as the eight sites surrounding it. When located on the same site or on neighbouring sites, two cells are in contact and each contact contributes an energy −*ε*. The adhesion energy of a cell at site *m* is therefore

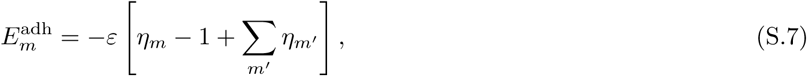

where the sum runs over all neighboring sites *m*^*′*^.

Each cell *l* consumes oxygen with an individual rate *Q*_*l*_ fixed by the local concentration *c*,

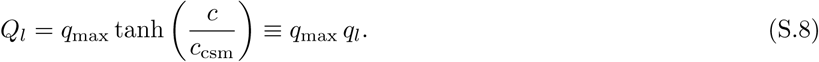

where *q*_max_ is the maximal consumption, *q*_*l*_ a rescaled consumption and *c*_csm_ a characteristic concentration. The cell consumption is almost constant above *c*_csm_ and approximately linear in *c* below *c*_csm_. The liquid film is not represented explicitly. Instead, the oxygen concentration field is evaluated from the cell positions by assuming it has reached a steady state. The method to do so relies on a Green function and is detailed below in Sec. G-2.

Each cell is endowed with aerotactic behavior, which sets in below a characteristic concentration *c*_aer_. We assume logarithmic sensing [G1]and a drift velocity *v*_aer_ ∼ ∇(ln *c*). To a move from site *m* to a neighbouring site *m′*, we associate an energy

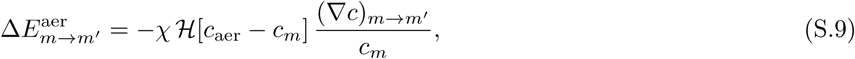

where *χ* quantifies the strength of aerotaxis, *c*_*m*_ is the oxygen concentration at site *m*, ℋ is a smoothed Heaviside function [G2], and (∇*c*)_*m→m′*_ is the concentration gradient at site *m* in the direction of site *m*^*′*^.

The motion of cells is simulated using a basic Monte Carlo (MC) algorithm. MC moves are only local, i.e. the cell move only to one of the neighbouring sites. The total energy change of a move from site *m* to site *m′* is

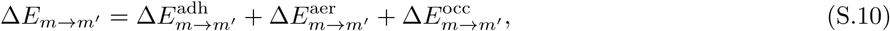

where 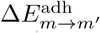 is the variation in adhesion energy. 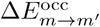 is infinite if the new site *m′* is already maximally occupied (*η*_*m′*_ = *η*_max_) and zero otherwise. A proposed move is accepted according to the Metropolis criterion, that is with probability

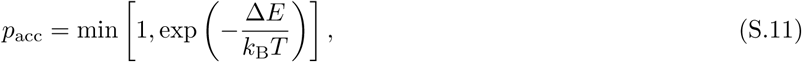

with *T* the temperature and *k*_B_ the Boltzmann constant. During a Monte Carlo step, each cell is subject to one proposed move.

As regards the units and parameters, the lattice spacing or cell size 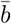 is taken as unit length and we fix 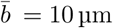. Accordingly, if each site is occupied on average by one cell, the dimensionless mean cell density is 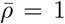, which corresponds to a real density 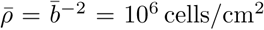. The energy unit is *k*_B_*T* and concentrations are made dimensionless with respect to the saturation value *c*_s_. The parameters of the model are listed in Tab. I, together with a default value which is justified below in Sec. G-3. Simulation units are understood when no unit is specified.

**TABLE I.** Units and parameters of the model. A default value, which holds unless indicated otherwise, is also indicated.

| symbol | value | quantity |
| --- | --- | --- |
| $\bar{b}$ | - | unit length, $\bar{b} \longleftrightarrow 10 \mu\text{m}$ |
| $k_B T$ | - | unit energy |
| $c_s$ | - | unit concentration |
| $\bar{\rho}$ | 1 | mean cell density |
| $\eta_{\max}$ | 4 | maximum occupation number |
| $\varepsilon$ | 0.15 | contact energy |
| $h$ | 1 | film height expressed in millimeter |
| $\kappa$ | 0.85/1.2 | field parameter as defined by Eq. (S.19) |
| $c_{\text{csm}}$ | 0.02 | concentration around which consumption becomes non-constant (Eq. (S.8)) |
| $c_{\text{aer}}$ | 0.1 | concentration below which aerotaxis sets in (Eq. (S.9)) |
| $\chi$ | 2 | aerotaxis strength (Eq. (S.9)) |
| $L$ | 100 | lattice size |
| $T_{\text{sim}}$ | $10^6$ | simulated time in MC steps |

## 2. Method

### 1. Green function

We consider a liquid film of height *h* whose top surface is in contact with air. Assume a (non-uniform) absorbing flux *j* is imposed at the bottom surface. What is the resulting oxygen concentration field in steady state? Here we give a partial response using a Green function. It is convenient to work with the shifted concentration

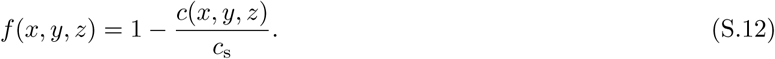

Choosing units here so that *h, D* and *c*_s_ are all unity, the equation and boundary conditions are

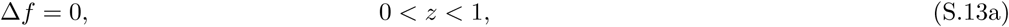

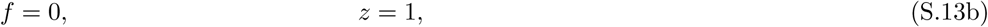

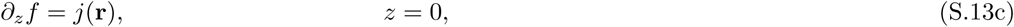

with Δ the Laplacian and **r** = (*x, y*) the position in the plane. Taking Fourier transforms with respect to *x* and *y* variables leads to

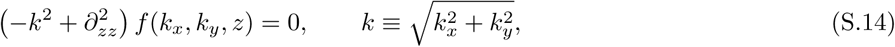

whose solution is a combination of exponential exp(±*kz*). Exploiting the boundary conditions gives

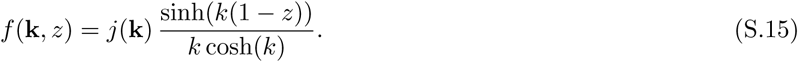

Considering only the surface and putting back dimensions, we get

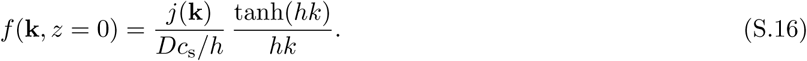

Note that this solution is valid only if the concentration remains positive everywhere.

Given a site **r**_*l*_ and a consumption *q*_*l*_ for each cell *l*, the local flux at a position **r** of the bottom substrate is

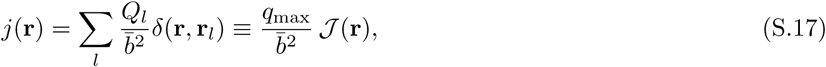

where *δ*(**r, r**_*l*_) is 1 when **r** belong to the site occupied by cell *l* and 0 otherwise, and the sum runs over all cells. The shifted concentration field is then

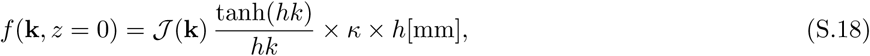

where *h*[mm] is the film thickness expressed in millimeter for convenience and *κ* a parameter defined as

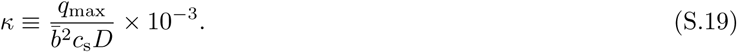

With *c*_s_ = 250 μM, *D* = 2 10^*−*5^ cm^2^ s^*−*1^ and *q*_max_ = 4.2 10^*−*17^ mol s^*−*1^, we have *κ* = 0.85, a value of order unity.

In practice, Fast Fourier transforms are used to switch between real and Fourier space representation of functions and obtain from Eq. (S.18) the shifted concentration at the bottom surface *f* (**r**, *z* = 0). The gradient of concentration is also computed in Fourier space.

### 2. Concentration field and cell consumption

The oxygen concentration field and individual cell consumptions are coupled variables. For any cell *l*, at any time, they should verify

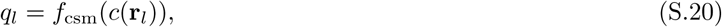

where the concentration *c*(**r**) depends on the whole set {*q*_*l*_} of cell consumptions. To ensure that this constraint holds, we used an iteration algorithm, which converges toward a fixed point solution of Eq. (S.20). We observe numerically that concentrations can be small, down to a few percent, but always remain positive as required.

In principle, the concentration field and cell consumptions should be updated each time a cell changes position. In practice however, this is exceedingly costly in computational resources as the concentration update is the limiting step in the simulation. Accordingly, we resort to a simple approximation where the field remains constant during one MC step and is updated after each MC step. Given that the MC moves considered are only local, that the oxygen concentration depends on many cells, and that aggregates evolve on time scale much longer than a single MC step, this approximation is well justified. As a side remark, we note that the transient decoupling between particles and field is similar to the methods used for simulations of block copolymers [G3].

### 3. Aggregate size

The size of aggregates is the main quantity of interest. In defining aggregates, only sites with occupation number *η* ⩾ 3 are considered. Using a connected component analysis, aggregates are identified and sorted in order of decreasing area 𝒜, defined as the number of sites composing the aggregate. We build a representative set by including aggregates, with larger ones coming first, until the sum of their areas exceeds 90% of the total number of aggregate sites. Such a definition allows to discard the small but potentially numerous aggregates and to focus on the larger aggregates that are of interest in this work. This definition is also suitable if there is a single or only a few aggregates. Finally, we compute the mean area ⟨𝒜⟩ of the aggregates in the representative set and use the radius 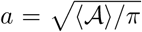 as a simple measure for the typical aggregate size.

### 3. Simulations

#### Choice of parameters

Here we briefly explain how the parameters were chosen. First, the typical dimension of *Dictyostelium discoideum* cells is fixed to 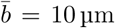, a convenient value that is approximately consistent with the range of cell size observed. Since the thickest aggregates reach 40 μm, as shown in SM-C, the maximum occupation number is set to *η*_max_ = 4. As regards the adhesion energy *ε*, and for simulations when only adhesion is present, we note the following. For *η*_max_*ε ≪* 1, aggregates are negligible in size. For *η*_max_*ε ≫*1, aggregation is irreversible: aggregates quickly form but remain frozen in size and do not coalesce further because cells are irreversibly stuck. The case *η*_max_*ε ≃* 1 is the interesting regime: large aggregates appear and coexist with a “gas” of isolated cells, with constant exchange between them. Small *ε* values favor a dense gas and aggregates with branched, irregular shapes while larger values yield more compact shapes but a gas of vanishing density. We choose *ε* = 0.15, in *k*_B_*T* units, as a compromise that features both rather rounded aggregates and a significant gas around them.

The typical concentration below which the individual consumption of cells becomes concentration-dependent is *c*_csm_ = 0.02, which is consistent with the range given by our measurements (see Fig. S4) Aerotactic behavior sets in below *c*_aer_ = 0.1 as found in Ref. [29]. The aerotactic strength is chosen so that when oxygen levels are everywhere below *c*_aer_, no sizeable aggregate form. A value *χ* = 2 is sufficient to enforce this requirement. Finally, the individual cell consumption was reduced by a factor 1.2, meaning that *κ* = 0.85*/*1.2 as indicated in Tab. I. This choice allows for a quantitative comparison between simulations and experiments. Note that a reduction in consumption may be physically understood because our two-dimensional model can not accurately describe the effects within three-dimensional aggregates, such as limited oxygen availability for cells most underneath.

As a side remark, we note that our model does not account for cell division. Even though our parameters may lead to oxygen concentrations above *c*_div_, the number of cells remains constant. Including cell division is far from straightforward because the very low concentrations involved (below *c*_div_) require the constant adjustment of individual cell consumption and drastically increase the computation time. This is why we have have chosen to focus on the interplay between adhesion, consumption and aerotaxis, which allows for an efficient exploration of parameter space and is sufficient to understand the mechanism underlying aggregates.

**FIG. S9.**
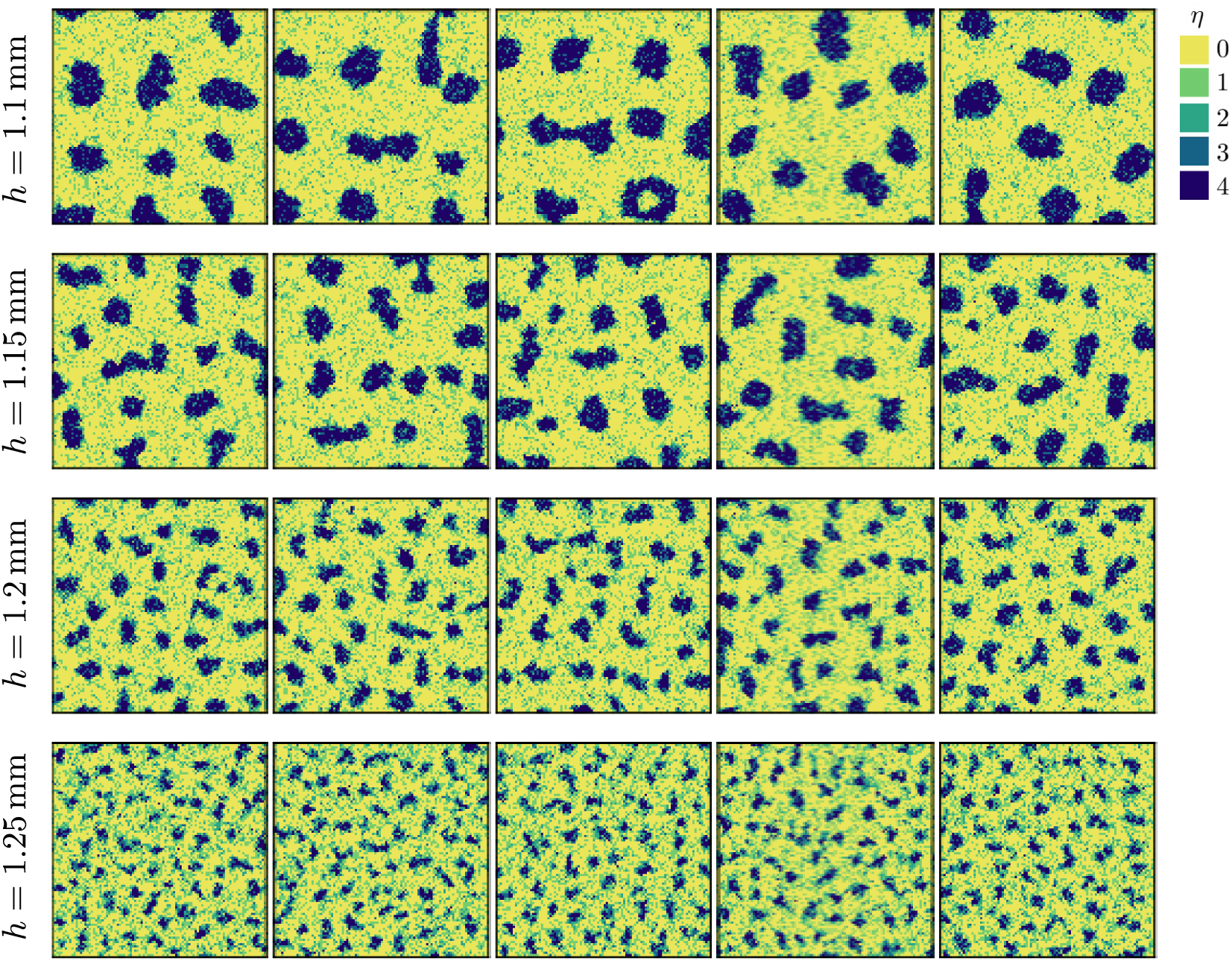
Aggregates in the cell-based simulations for various film height *h*. The average density is 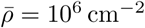, the system size 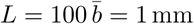, and the simulations involve 10^4^ cells. From top to bottom, the aggregates have typical size *a* = 90, 65, 42, 26±5 μm. From left to right, the simulation time is *t* = 5, 6, 7, 8, 9 × 10^6^ MC steps. The aggregates have reached a steady state but never settle into a static configuration.

#### Shape of aggregates

A series of aggregate configurations are shown in Fig. S9. As regards the shape aggregate, several comments are in order. First, simulated aggregates are not always quasi-circular but can transiently adopt elongated shapes that are rarely seen in experiments. Second, the “gas” of cells outside the domains is rather homogeneous. This is somewhat different from experiments (see Fig. 1) where the largest aggregates are surrounded by domains that are smaller and of lower density. These two differences indicate that our simulations provide only an idealized description of the phenomenon. Third, an aggregate can occasionally be seen with a hole, at least transiently. When too large with respect to its equilibrium size, a domain may nucleate a hole in its center and subsequently break into smaller pieces. While the hole is presumably an artifact of our two-dimensional description, it points to the aggregate interior as the location where the aerotactic effects are the strongest.

#### Steady state

For each film height, two initial conditions were considered: *(i)* cells are placed at random, or *(ii)* cells are arranged on a single giant aggregate of circular shape within which *η* = *η*_max_. Though the relaxation may require a significant time – several millions of MC steps for the largest aggregates –, the typical aggregate size eventually reaches a plateau independent of the initial conditions, which indicates that a steady state has been attained. Nevertheless, it is clear from Fig. S9 that the aggregates never settle in a static configuration (see also movie M6). Domains continually deform and rearrange, without any trend to further coarsening. This observation shows that domain growth is not kinetically limited and that a genuine steady state with a preferred domain size has been reached. Such a dynamic behavior of domains is also seen in experiments (SM-A).

## H. MOVIES

### Movie M1: Microphase separation in *Dictyostelium discoideum* under submerged conditions

Cells were plated at *t* = 0 at 7.5 10^4^ cell/cm^2^ (not shown). Cells first grow exponentially and at time *t* = 24 h (first image of the movie), the density is close to 4.5 10^5^ cell/cm^2^. At *t* = 35 h, when the density reached 7.5 10^5^ cell/cm^2^, cells start to gather on the right side, forming small transient loose aggregates. At *t* = 60 h, the full field of view is invaded by aggregates. These aggregates grow in a very dynamic way: they exchange single cells with the surrounding and fuse with other clusters to become bigger. Occasionally, they fully melt or divide in a fission event. Between 60 h and 114 h, almost 2.5 days, they move continuously but never agglomerate into a giant aggregate. Microscope settings: X4 objective lens, confocal in transmission mode. Time label is in hours. Bar is 500 μm.

### Movie M2: Higher magnification, one hour sequence, of an aggregate 2.5 days after entering in the microphase-separated state

The aggregate lies on a first carpet of flattened single cells in close contact with the substrate. These flattened cells on the substrate layer are poorly mobile and still isolated; they do not form a dense monolayer. The aggregate is moving by exchanging a few cells with the gas phase but it keeps most of its constituent cells as well, displaying large deformation and apparent traction from its periphery. The focus plane on this movie is around the background of cells and the inner part of the aggregate appears partially transparent. Microscope settings: X10 objective lens, confocal in transmission mode. Time label is min:sec, bar is 50 μm.

### Movie M3: Effect of EDTA on aggregate size

Aggregates were formed for 2.5 days in normoxic conditions (21% O_2_). The EDTA was added at time *t* = 0, which is the first frame of the movie. Microscope settings: X4 objective, inverted microscope in transmission mode. Time label is h:min, bar is 1000 μm.

### Movie M4: Dynamics of aggregates in the steady microphase-separated state

Large field of view (8.33 × 6.25 mm^2^) of the aggregate displacements between *t* = 40 h and *t* = 64 h after plating the cells. The medium height is *h* = 1.5 mm. Microscope settings: X1 objective lens, Leica MZ16 binocular equipped with a TL3000 Ergo transmitted LED light base. Time label is min:sec, bar is 50 μm.

### Movie M5: Effect of changing the atmosphere oxygen level on aggregate size

Aggregates were formed for 2.5 days in normoxic conditions (∼21% O_2_) before taking the first image. Right after the first image referred as *t* = 0, the oxygen level was decreased to 10% by injecting N_2_, before returning to 21% at *t* =1:40 (h:min). After the descending step, the aggregates slowly shrink and break in small aggregates. After the ascending step, the aggregates slowly regrow. Microscope settings: X4 objective, inverted microscope in transmission mode. Time label is h:min, bar is 200 μm.

### Movie M6: Simulated aggregates in steady state

The system has size 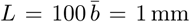, density 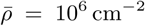 and includes 10^4^ cells. With film height *h* = 1.15 mm, the typical aggregate size is *a* = 65 μm. To reach steady state, 5 10^6^ MC steps were performed before the movie starts. The total duration shown is *T* = 2 10^5^ MC steps.

